# Sleep-state dependent cerebellar processing in adult mice

**DOI:** 10.1101/2023.10.30.564769

**Authors:** Cathrin B. Canto, Staf Bauer, Tycho M. Hoogland, Hugo H. Hoedemaker, Cynthia Geelen, Sebastian Loyola, Pablo Miaja, Chris I. De Zeeuw

**Affiliations:** Department of Cerebellar Coordination and Cognition, Netherlands Institute for Neuroscience, Royal Netherlands Academy of Arts and Sciences, Amsterdam, The Netherlands; Department of Neuroscience, Erasmus MC, Rotterdam, The Netherlands

## Abstract

The cerebellum is important for motor performance and adaptation as well as cognition. Sleep is essential for optimizing of all these functions, but it remains to be elucidated how sleep affects cerebellar processing. It has been suggested that sleep periods with muscle twitches entrain the cerebellum with a copy of motor commands and subsequent sensory feedback signals, to develop predictive coding of movements. If this hypothesis is correct, one expects phasic correlations between the muscle twitches and specific features of the electro-encephalography (EEG) recordings in the cerebellum during various sleep stages as well as the climbing fiber activity in the cerebellar cortex, the modulation of which is relayed from the cerebral cortex via mesodiencephalic junction and inferior olive.

Here we provide evidence for coherent correlations between cerebellar and cerebral cortical sleep spindles, twitches as well as patterns of climbing fiber activity. Our data are compatible with the novel concept that muscle twitches evoke complex spike synchronicity during NREM, which in turn affects cerebellar spindle activity and cerebellar-cortical information flow, thereby entraining an internal forward model.

## Introduction

Perception and action depend on the state of the embodied self. How sensory and motor inputs are perceived and how this in turn leads to a well-timed and accurate motor action and motor memory formation relies on the cerebellum. Indeed, the cerebellum is a brain region known for its role in motor control, planning and coordination of voluntary and fine movements, muscle tone, balance and posture, as well as non-motor functions, such as cognition^1–6^. The anatomical reasoning behind such complex functionality is that the cerebellum receives converging direct inputs from multiple parts of the brainstem, while it has extensive reciprocal connections, albeit more indirectly, with the frontal, parietal and temporal cortices^7^. In the context of motor control, the cerebellum receives two major inputs, climbing fiber activity, the modulation of which is relayed from the cerebral cortex via mesodiencephalic junction and inferior olive, and mossy fiber activity that transfers motor information received by the pontine nuclei, to estimate the future direction of movement, in line with the operation of a forward model ^8–11^. As a consequence, the cerebellum and thereby also the related control system of the embodied self can predict how different levels of actions may alter sensory inputs and events^11–13^. These predictions work optimally when the forward model is frequently calibrated through error-based learning.

Recently, it has been proposed that sensory feedback of sleep-dependent, subcortically generated myoclonic twitches of the limbs promote development of the functional connectivity between the cerebellum and its input and output structures^14,15^. In general, sleep has been shown to be vital, supporting essential physiological and psychological functions, but the specific role of sleep for the functionality of the adult cerebellum is still an almost uncharted territory^16^. Given that sleep after cerebellum-dependent eyeblink conditioning ^17–20^ improves memory consolidation and retrieval in mice as well as humans ^21–24^, we hypothesize that sleep plays a significant role in subconscious cerebellar (implicit) learning, through opening a gate for cerebellar to cortical sensory transmission ^25–27^ and through replay mechanism similarly to those that have been shown for conscious hippocampal and cerebral (explicit) learning ^28–34^. Indeed, neuronal oscillations and memory reactivation in these latter regions have been shown to contribute to improved declarative memory performance in various conditions ^28–33^. If our hypothesis on the role of sleep in cerebellar learning is correct, one expects temporal correlations between the muscle twitches and specific features of the electro-encephalography (EEG) recordings in the cerebellum during various sleep stages. Moreover, if correct, one also expects phasic correlations with the so-called climbing fiber activity in the cerebellar cortex, the modulation of which is readily relayed from the cerebral cortex and mesodiencephalic junction to the source of the climbing fibers, the inferior olive ^35^. In contrast, the modulations of the mossy fiber system may be more subject to downstate mechanisms during sleep ^36^.

To test this hypothesis, we set out to record the different sleep stages with local field potentials (LFPs)/electroencephalogram (EEG), electro-oculography (EOG) and electromyography (EMG) ^37^ in adult mice. We divided the non-rapid eye movement (NREM) sleep into NREM1-3, with sleep spindles and slow waves in the cerebral cortex mainly occurring in NREM2 and NREM3, respectively ^38,39^. Rapid-eye movement (REM) sleep is associated with dreaming ^40,41^, and was characterized by theta activity occurring in the hippocampus (5-8Hz in rodents) ^42–44^. In short, our data provides evidence that the cerebellum shows sleep-state dependent activity ^45–48,^. More specifically, we found NREM sleep-dependent cerebellar activity with signals having a lower amplitude compared to local field potentials (LFPs) in the cerebral cortex, and with cerebellar spindles and slow-waves associated with NREM2/3 and NREM3, respectively. In general, the cerebellar sleep spindles in adult mice correlated with cortical spindles, and the cerebellar spindle activity co-occurred with twitches^49,50^. Moreover, exploiting Ca^2+^ imaging of unrestrained sleeping animals with the use of a miniature microscope ^51^, we revealed that twitches during NREM stages evoked climbing fiber induced calcium transients, akin complex spikes, in the cerebellar cortex and increased complex spike co-activation. Synchronization may open the gate for cerebello-cortical information transfer. We find evidence for this in our LFP/EEG recordings using directed-coherence ^48,52,53^ to uncover information transfer from the cerebellum to the cortex. Our data suggest functional connectivity among distant sensorimotor structures, including cerebellar and cerebral cortex, increases during NREM spindle activity and twitching. To the best of our knowledge, our data present the first evidence for a mechanism of how sleep may gate to improve long-range functional connectivity, thereby bridging memory formation and consolidation across the declarative and procedural domains ^15,27,54,55^.

## Results

We first assessed to what extend the cerebellum displays sleep-state dependent activity by recording during waking and natural sleep using polysomnographic recordings in unrestrained mice (n=8), The cerebellum displayed NREM activity, resembling cortical activity (**Figure 1A**). Local field potentials were obtained from electrodes implanted in lobule IV and V of the cerebellar cortex and in the ipsi- and contralateral primary motor cortex. The cerebellum showed NREM2-related sleep spindles (9-16 Hz) with highest spindle power during NREM2 and slow-waves with highest slow-wave power (0.5-4Hz) during slow wave sleep, similar to what has been previously described in non-human primates and rats ^46–48^. However, slow-wave sleep in the cerebellum had a lower amplitude compared to the cortex **(Figure 1A)**. Thus, the mouse cerebellum shows sleep-dependent NREM activity with sleep spindle, and slow-wave sleep-dependent oscillations.

**Figure 1:**
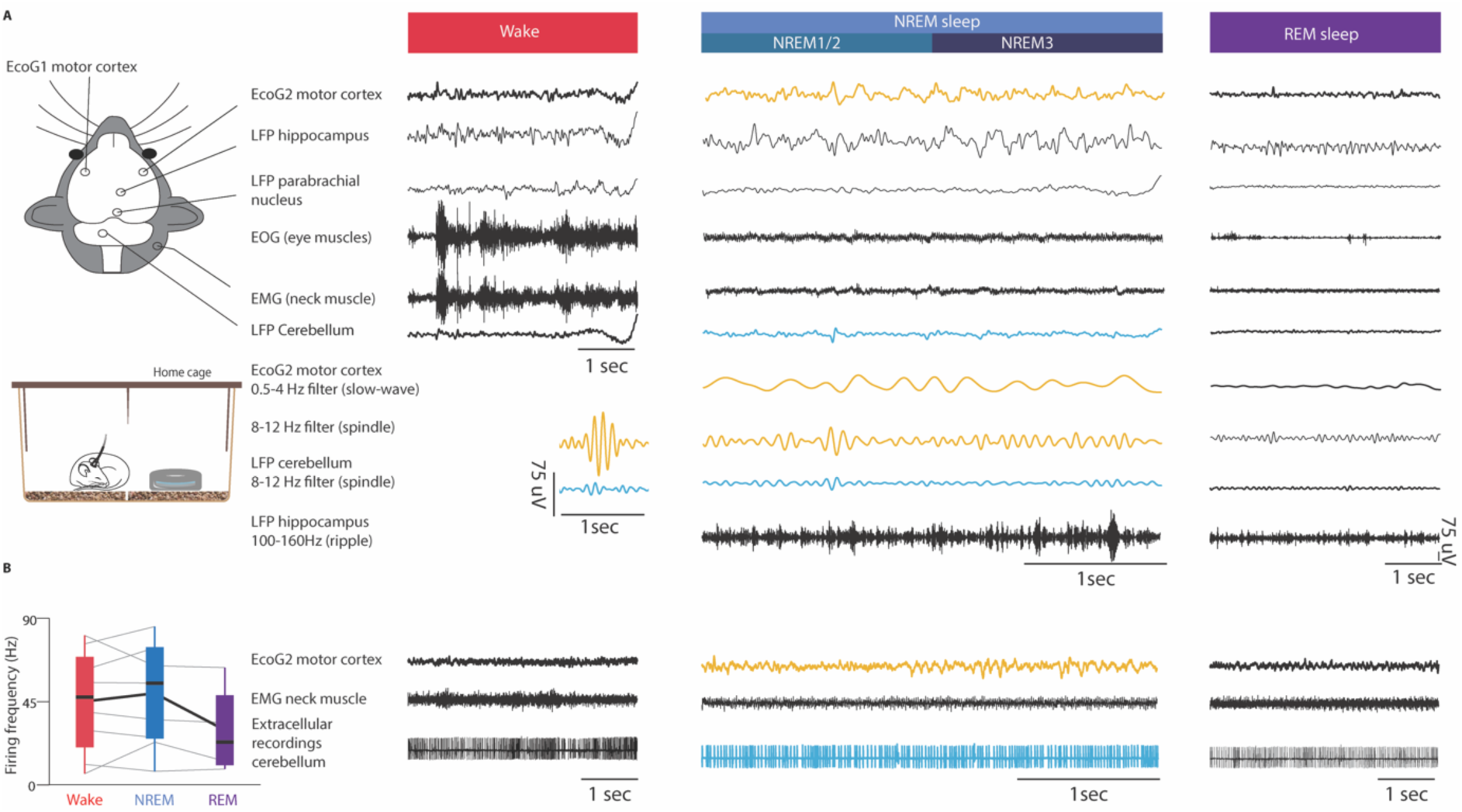
Seep-state dependent activity in the cerebellum. A) NREM sleep-related cerebellar spindles (NREM2, light blue) and slow waves (NREM3, dark blue) are present in cerebellum, but with lower amplitude compared to the cortex. REM sleep (purple) was defined by theta frequency activity in the hippocampus. We performed electrocorticographical (EcoG) recordings of the motor cortex, local field potentials (LFPs) of the hippocampus, parabrachial nucleus and cerebellum, electrooculographical (EOG) and electromyographical (EMG) recordings of the neck muscle in unrestrained animals. Animals were placed in their home cages from 1 up to 24 hours. For visualization purposes we also filtered the data to show spindle and ripple activity in the cortex (yellow) and cerebellum (light blue) as well as the hippocampus (black), respectively. B) Individual Purkinje cells of the cerebellar cortex, recorded extracellularly in-vivo in head-fixed sleeping mice, show either increases or decreases in firing frequency (left) during sleep compared to wakefulness but not all neurons show a similar response. Right, example of one Purkinje cell recorded extracellularly during wakefulness, NREM and REM sleep.

Next, we recorded the extracellular activity of individual cerebellar neurons in head-fixed mice during wakefulness and sleep (**Figure 1B**). The climbing fibers and mossy fiber–parallel fiber system converge on Purkinje cells and cerebellar nuclei neurons, which form the output of the cerebellar cortex and cerebellum as a whole, respectively. Activity in climbing fibers elicits so-called complex spikes in Purkinje cells, that of mossy fibers and parallel fibers modulates Purkinje cells’ simple spike activity. Individual Purkinje cells. Individual neurons show sleep-related modulation in their simple spike firing frequency, however not all neurons modulate in a similar way. Three out of 8 neurons showed a decrease in simple spike firing during NREM sleep, whereas 3 neurons showed an increase in the simple spike firing frequency during NREM (**Figure 1B)**.

### Cerebellar sleep spindles in adult mice

The presence of spindles in the cerebellum is intriguing and was first described in non-human primates^48^. We isolated spindles in the cortex and cerebellum (**Figure 1A**) to assess whether these spindles have similar frequencies as the thalamo-cortical spindles with similar spindle duration (**Figures 2A-2C**). Cerebellar spindles indeed have similar frequencies as the thalamo-cortical spindles in adult mice, ranging on average from 11.3-12.0 Hz, with a mean and median of 11.67 Hz and 11.7 Hz, respectively, compared to cortical spindles having spindles in the frequency ranging on average from 11.2-12.0 Hz, with a mean and median of 11.8 Hz and 11.9 Hz, respectively (**Figure 2B**, p = 0.23, paired t-test n.s). The duration of the cerebellar spindles lasted from 0.87-1.03 s which matches cortical spindles that last from 0.83-1.03 s (**Figure 2C**, p = 0.43, n.s.). Analyzing phase preferences of individual Purkinje neurons of the cerebellar cortex, revealed that not all neurons show a phase preference with respect to spindle frequencies during sleep but some do (3 out of 13 neurons show significant z-score >2.5, phase preference lies in the first part of the rising phase). In human infants it has been shown that spindles correlate with myoclonic twitches of limbs during NREM sleep^49^. Here we wanted to determine if so-called myoclonic muscle twitches occur in adult mice during NREM sleep, and if so, whether they correlate with both cerebellar and cortical spindles.

**Figure 2:**
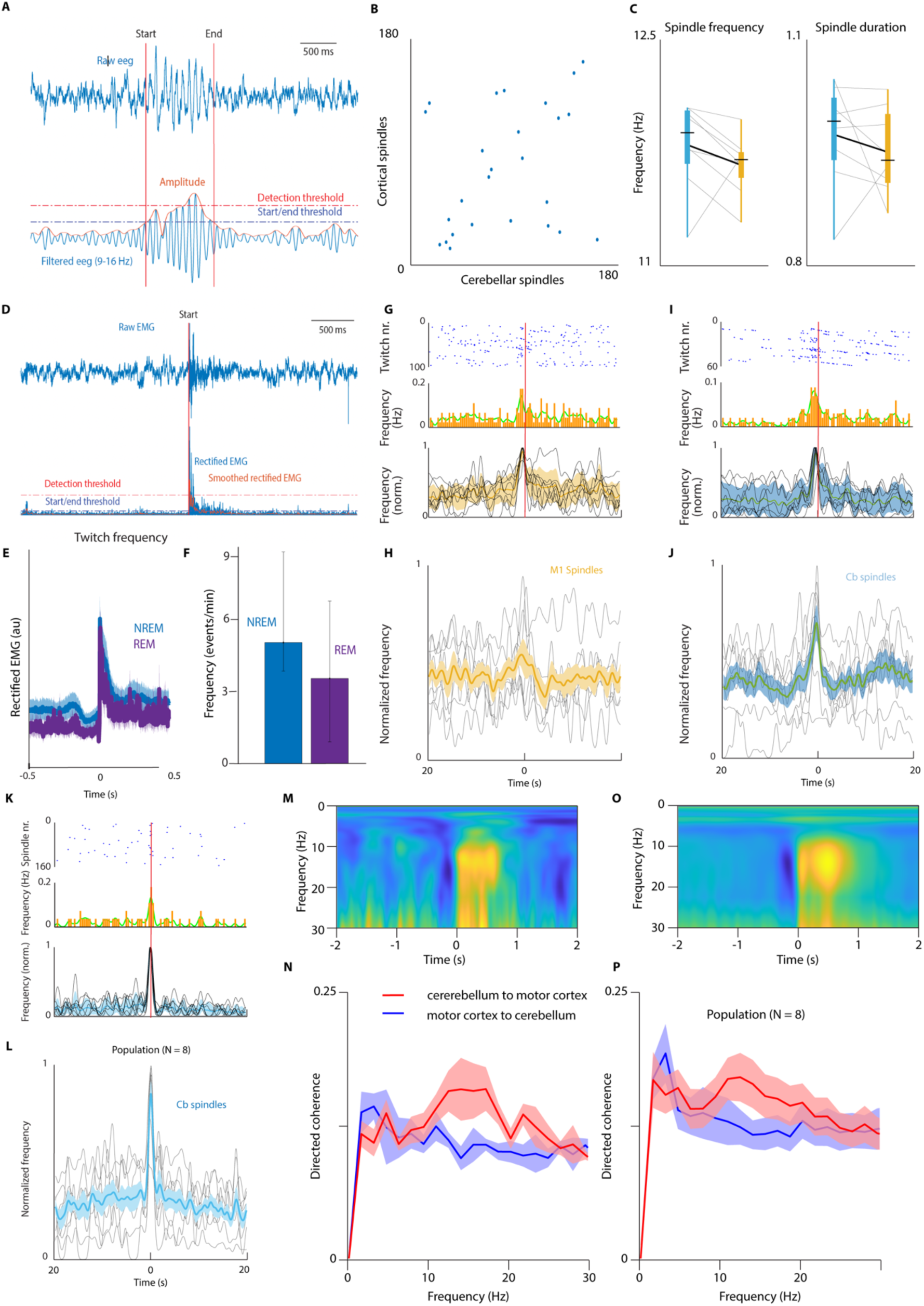
NREM sleep-related muscle twitches co-occur with cerebellar sleep spindles, associated with directional information flow from the cerebellum to the cortex. A) To analyze sleep spindles, we filtered data between 9-16 Hz, set a threshold and ran through different wake and sleep stages. B) We plotted cerebellar spindles against the spindles in the motor cortex and compared C) the frequency and duration for the cerebellum and cortex. D) To analyze muscle twitches, we took the smoothened rectified EMG, calculated a detection and start/end threshold based on the mean standard deviation and searched for twitches. E) The rectified EMG and F) twitch frequency compared between NREM (blue) and REM (purple) sleep revealed a similar twitch and frequency of twitches during NREM and REM sleep. G,I,K) A scatterplot (top), frequency plot (middle) and normalized frequency plot (bottom) showing spindles (G,I, K) at time interval before and after onset of twitches (G,I, H and J), or K, L) cortical spindle activity. Twitches co-occur with cerebellar spindles and cerebellar spindles correlate with cortical spindles. M, O) Power frequency plots and directed coherence plots (N, P) revealing spindle activity at a single mouse level (M, N) versus population activity of 8 mice (O, P). Directed coherence reveals cerebellar to cortical communication during spindle frequencies.

### Muscle twitches during REM and NREM sleep in adult mice

Performing electromyographic recordings of the neck muscles in adult mice revealed muscle twitches during NREM and REM sleep (**Figures 2D and 2E**). Those muscle twitches were isolated using the same method as used for developmental muscle twitches. Those twitches occurred both during NREM and REM sleep (**Figures 2E and 2F**). Although we did not record from limb muscles, we demonstrate that neck muscles twitch during NREM sleep in adult mice. We could thus examine whether NREM sleep related muscle twitches correlate with cerebellar spindles.

### Muscle twitches are associated with cerebellar sleep spindles

Similar correlations are observed between muscle twitches and cortical spindles in human infants. In this study, we present evidence that in adult mice, cortical spindles also manifest in conjunction with muscle twitches, as depicted in Figure 2G and 2H, though the association is less pronounced than with cerebellar spindles (shown in **Figures 2I and 2J**, and further highlighted in the scatter plots above **Figures 2G and 2I**). The correlation of muscle twitches with cerebellar spindle activity in adult mice implies that muscle twitches during NREM sleep are not arbitrary or without function. Instead, they may play a significant role in facilitating information transfer between the cerebellum and the cerebral cortex.

### Cerebellar to cortical communication during sleep-dependent spindle frequencies

It has been proposed that spindles are maintained through thalamo-thalamic or thalamo-cortical interactions, with their waves of activity moving at the same speed of direct neuronal communication ^56,57^. First, we analyzed whether cerebellar spindles occur around cortical spindles (**Figures 2K and 2L**). We then determined the directionally of the interactions between the cerebellum and the cortex, using directed coherence spectra. Directed coherence at spindle frequencies revealed a directionality from the cerebellum to the neocortex, at the single mouse level (**Figures 2M and 2N**) as well as population level (**Figures 2O and 2P**). These data suggest that information flows from the cerebellum to the motor cortex during spindle frequencies, with directed coherence being significantly greater (p<0.05). Given that twitches occur around spindle activity, and that spindles are associated with directed information flow from the cerebellum to the cortex, we were interested in the mechanisms underlying this sort of sleep-dependent encoding during NREM sleep.

### Twitch evoked synchronized complex spike activity in Purkinje cell dendrites

Finding twitch and cerebellar spindle correlations motivated us to study whether the sensory feedback from muscle activity leads to complex spike firing in Purkinje cells (PCs), which are the sole output neurons of the cerebellar cortex (**Figure 3**). Single neuron analysis cannot reveal PC population activity, which needs to be analyzed to show how twitch evoked sensory feedback enters the cerebellum. NREM sleep has been suggested to play a role in cognition ^38,56,58–69^. We hypothesize that if cerebellar sleep-related activity supports such a role, not all PC dendrites will receive the same information during sleep with co-activation becoming sparser, and by this sensory feedback becomes more specific. Therefore, we performed miniscope imaging from populations of PC dendrites in lobule IV/V of the cerebellum in conjunction with cortical EcoG, hippocampal LFP and EMG recordings of the neck (N=9 mice) (**Figures 3B and 3C**). Across our recordings mice spent most time in NREM2 (49,4%) followed by NREM3 (20.1%), and REM (5.3%) sleep. Thus, the highest fraction of scored sleep was spent in the NREM2 stage (NREM1: 6.5%, NREM2: 60.1%, NREM3: 24.5%, REM: 8.9 %). During the transition from wake (Wake: 0.95±0.6 Hz, 1363 PC dendrites) to NREM1 sleep (1.0±0.5 Hz, 1277 PC dendrites) complex spike firing rates showed a small but significant increase (**Figure 3D**). Complex spike rates dropped significantly from NREM1 to NREM2 (0.77±0.5 Hz, 1836 PC dendrites) and NREM3 (0.81±0.5 Hz, 1418 PC dendrites) sleep. Complex spike rates increased to levels indistinguishable from wakefulness during REM sleep (0.94±0.5 Hz, 1498 PC dendrites). Significance was confirmed using a Kruskal-Wallis test comparing among groups followed by a Scheffe’s procedure for multiple comparisons (ɑ=0.01). Historically the complex spike has been seen as an all-or-none event, where the climbing fiber generates a massive synaptic response. Recent work has cast this idea aside suggesting that varying levels of sensory input to the inferior olive, which gives rise to the climbing fibers, could generate more graded complex spike responses ^70,71^, as reflected potentially in a variable number of spikelet waves associated with a complex spike. Calcium imaging revealed that a graded response was reflected in the amplitude of complex spike evoked calcium transients^71^. We reasoned that if sensory input is gated during sleep, this should be associated with reduced complex spike -evoked calcium transient amplitudes. Indeed, relative to wakefulness (Wake: 12.4±9.7, 3972 transients, expressed as peak amplitudes, a denoised and scaled version of ΔF, mean±SD) the calcium transient amplitudes were reduced (**Figure 3E**) across all sleep states (NREM1: 11.1±10, 5517 transients; NREM2:10.8±7.7, 6082 transients; NREM3: 10±7.3, 5429 transients; REM: 10.6±8.2, 6272 transients). Inter-group differences were significant for Wake-NREM1, Wake-NREM2, Wake-NREM3, Wake-REM, NREM1-NREM2, NREM1-NREM3, NREM2-REM and NREM3-REM (Kruskal-Wallis test with post-hoc Scheffe test with ɑ=0.01). Given that our imaging approach allowed monitoring of multiple Purkinje cell dendrites we asked whether the rate at which at least 10% of all detected Purkinje cells co-fired within the imaging field-of-view changed during sleep (**Figure 3F**). There was a significant drop in co-activation rates from wakefulness (2.87±1.5 Hz) to NREM and REM (NREM1:1.5±1.8 Hz; NREM2: 1.0±1.2 Hz; NREM3: 0.77±1.2 Hz; REM: 0.96±1.4 Hz) sleep relative to wakefulness (Kruskal-Wallis test with post-hoc Scheffe test with ɑ=0.01). The co-activation rate dropped significantly from NREM1 to NREM2 and NREM3 sleep. Thus, even though there were several epochs of co-activation encompassing more than 10% of Purkinje cells during sleep they became sparser.

**Figure 3:**
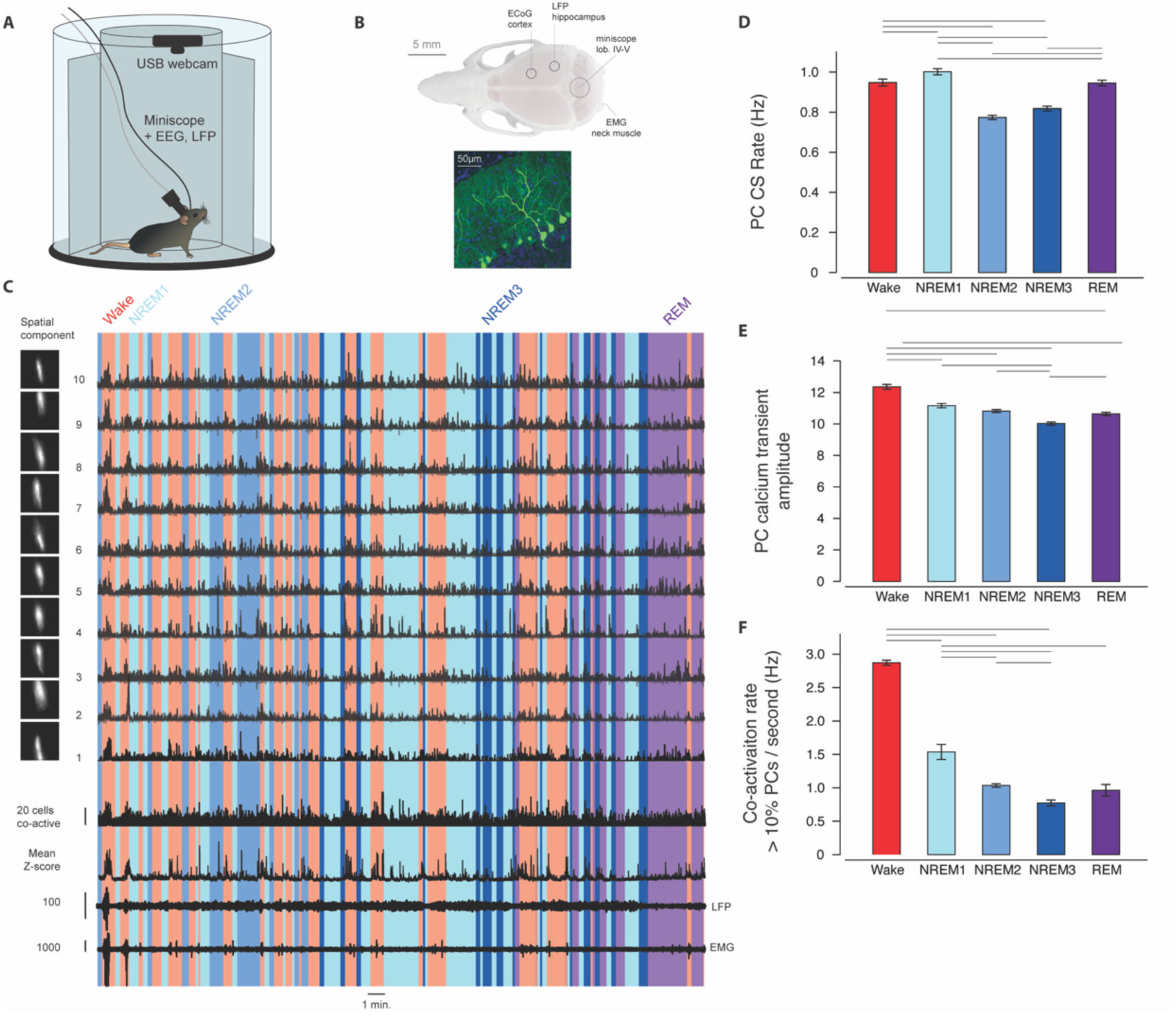
Co-active complex spike activity in Purkinje cell dendrites during wakefulness and sleep. A) To increase the chance of spontaneous sleep, C57/BL6 mice were placed in a custom modified slowly rotating drum for a period of up to four hours during which the movement kept the animals awake. After this period animals were allowed to fall asleep either in a non-moving drum with bedding material from the home cage or in the home cage. B) Both cortical electroencephalogram (EEG) and hippocampal local field potential (LFP) were recorded to monitor wake and sleep (top). B) Purkinje cell dendrites of cerebellar lobule IV/V were selectively transduced with the genetically encoded calcium indicator GCaMP6f (bottom). C) Calcium transients were recorded from individual PC dendrites that had large amplitude, fast onset times and kinetics separating them from other sources of calcium entry ^72^. Such transients were clearly resolvable and allowed us to extract complex spike onset times after signal deconvolution. All data was passed through motion correction using NorMCorre ^73^ followed by signal extraction using CNMF-E as described previously ^51^. Cellular resolution imaging of PC dendrites allowed us to assess CS activity in dozens of cells at once while preserving the topology of the individual neurons with a field-of-view of 785 by 502 µm. Using our data, we extracted 74±21.6 PC dendrites on average (range: 44-119, N=9 animals). A widely used standard in the sleep field is to sample epochs lasting four seconds and score them into sleep states. We adapted this approach for this study by scoring sleep into four second epochs and subsequently performing analyses on complex spike evoked calcium transient activity within them. Colors indicate wakefulness (red), NREM1 (light blue), NREM2 (blue) and NREM3 (dark blue) as well as REM sleep (purple). The spatial footprints of 10 Purkinje cell dendrites are shown (left column) together with the extracted signal for these dendrites over a period lasting several minutes. All signals were deconvoluted and events representing the onset of a transient were selected. Occurrence of complex spike (CS) events were summed to obtain the co-activation of CS associated calcium transients in PCs. Modulation of such co-activation, or the mean Z-scored signal were apparent during clear movements picked up in the EMG channel (units for LFP and EMG represent uV). D, E, F) Quantification of complex spike activity imaged during sleep. D) complex spike (CS) rates across sleep stages E) PC calcium transient amplitudes, and F) the co-activation rate of >10% PCs / second (Hz) was plotted for Purkinje cells (PCs). Lines above bar graphs indicate significance between groups established using a Sheffe’s test for multiple comparison. Error bars represent standard error of the mean.

Next, we wanted to assess whether twitches lead to complex spike co-activation around twitch onset as a potential result of sensory feedback from activated muscles. Therefore, we scored the neck muscle EMG for brief high frequency deflections reflecting a muscle twitch and used these as onsets to determine if complex spike-evoked calcium transients could be triggered off of these twitches (**Figure 4**). The sharp breaks in the rectified EMGs at t=0 highlight the onset of the twitches. The largest fraction of twitches occurred during wakefulness (1.2 Hz, twitches per second). The rate was significantly suppressed, but present in both NREM and REM stages (NREM1: 0.4 Hz, NREM2: 0.5 Hz, NREM3: 0.3 Hz, REM: 0.3 Hz). During wakefulness and across all sleep stages, twitches were associated with complex spike-evoked calcium transients that peaked after twitch onset. However, the onset of CS activity was shortly prior to the onset of the twitch which may indicate that the cerebellum receives a motor command copy of the twitch prior to the sensory feedback. During REM a pronounced double peak response was found after twitch onset supporting the idea that twitches during NREM and REM sleep might fulfill a different functionality but both leading to an increase in CS activity, likely allowing inputs to reach threshold and information transfer from the cerebellum to the CN and beyond ^74^.

**Figure 4:**
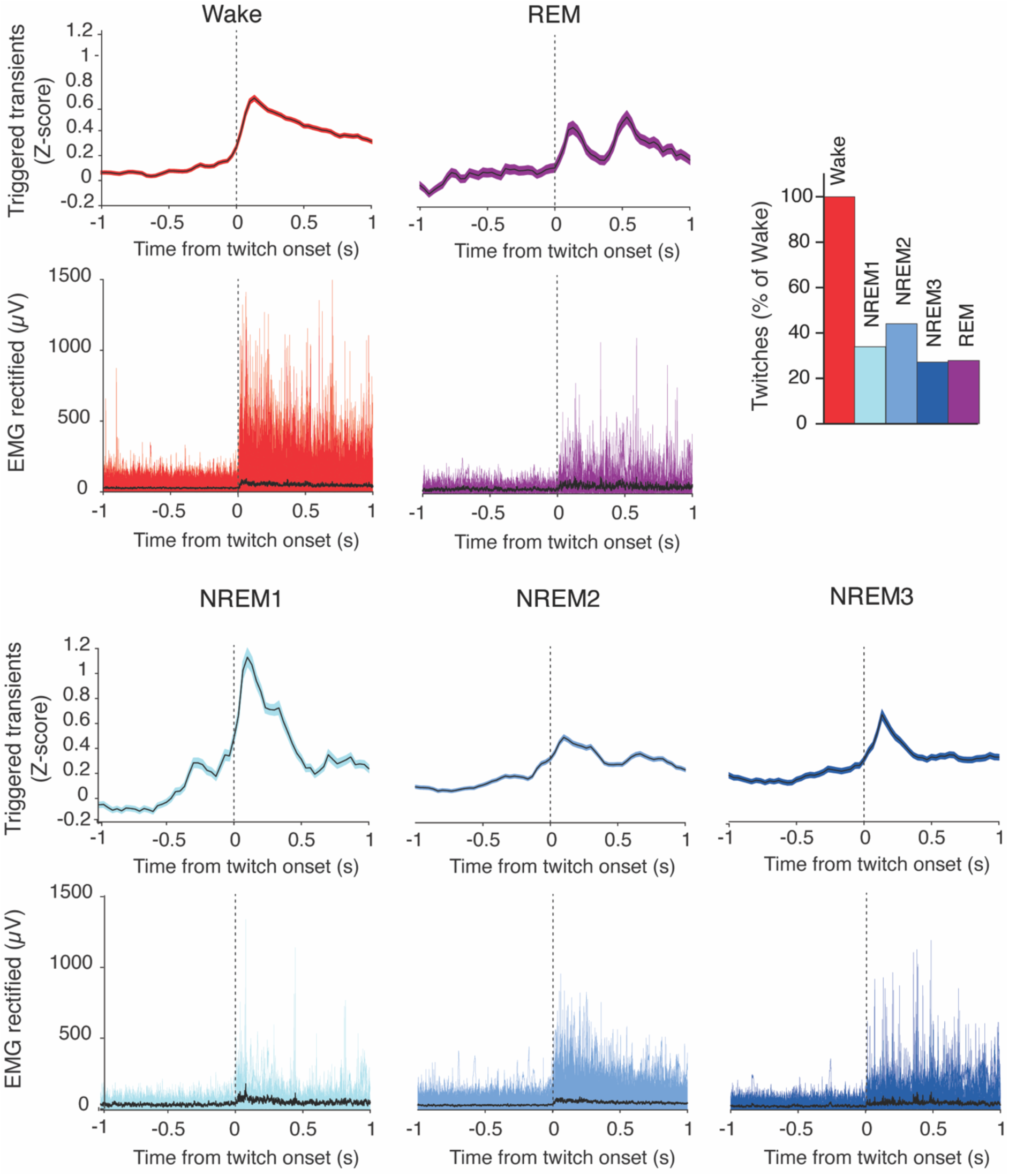
Twitch onset-triggered complex spike evoked calcium transients during sleep. Complex spikes evoked calcium transients in Purkinje cell dendrites (top) were triggered off the onset of twitches detected in neck muscle EMGs (bottom showing rectified EMG). Most twitches occurred during wakefulness and their number dropped substantially across all sleep states. Peak responses occurred after onset of the movement. Shaded areas in top panels represent standard error of the mean.

## Discussion

Sleep is important for the consolidation and retrieval of cerebellar-dependent motor memories in adult mice and such learning is facilitated by extending the permitted sleep period after cerebellar motor learning ^21^. We present data here that provides insight into the physiological changes that occur in the cerebellum at the cellular and system level to store such memories.

### Sleep-related cerebellar activity revealed with polysomnographic and miniscope recordings

Our data demonstrate sleep-related cerebellar activity in adult mice, with waxing and waning oscillations in the spindle frequency band (9-16 Hz) during NREM2 and NREM3 and slow waves during NREM3. These data recapitulate findings in non-human primates, where spindle oscillations were recorded during sleep as well as previously recorded slow wave activity in mice ^46,47^. Moreover, we demonstrate that Purkinje cells show different sleep modulation, with the simple spike modulation being inconsistent across all neurons recorded, similar to what has been shown in mice before ^45^. In sleeping cats and macaques there is an increase in Purkinje cell firing during REM sleep ^75–77^.The differences in sleep modulation of simple spikes in our data and recordings in other species may lie in the recording location and the associated differences in baseline properties of individual neurons ^78,79^. Complex spike rates, recorded with miniature microscopes, are depressed during NREM2 and NREM3 sleep. By contrast the complex spike rates during NREM1 sleep were slightly elevated relative to wakefulness and REM sleep. Up to now the literature on complex spike activity is contradictory. For example, recordings from cat cerebellum revealed modulation during synchronized (NREM) and desynchronized (REM) sleep states in Purkinje cell simple and complex spiking ^80^. Complex spike firing during REM sleep in cat increased in the absence of eye movement and decreased when eye movements were present. A contrasting study in macaques failed to find such a distinction ^75^ and instead found a reduced rate of complex firing during both NREM and REM stages of sleep in a select set of eight PCs. Later follow up studies in cat cerebellum revealed mostly a suppression of simple spikes and complex spikes during NREM stages of sleep and an increase during REM sleep that was partially attributable to eye movements. A recent study presented the first cerebellar electrophysiological recordings during sleep in mice ^45^. PC activity displayed an overall decrease during both NREM and REM sleep. Moreover, none of the aforementioned work has clearly distinguished modulation of cerebellar PC activity across the different stages of NREM sleep. Using miniscope imaging in unrestrained mice, we now show that complex spike activity in ensembles of PCs in the anterior vermis is modulated across the different sleep stages both in their rate, amplitude, and levels of co-activation. Whether these data can be extrapolated to e.g., the posterior cerebellar lobe needs to be studied but a reduction and refinement in complex spike activity suggests that activation is more sparse during sleep, supporting sleep-specific synergistic plasticity in the cerebellum leading to specific synapses to be formed or lost ^81^.

### Reduced amplitudes of complex spikes during NREM and REM sleep

Climbing fibers have typically been considered to trigger all-or-none complex spike responses in Purkinje cells. However, recent calcium imaging of sensory evoked complex spikes have challenged this assumption and suggest that a more graded response can be elicited by modulating the complex spike waveform and its number of spikelets ^71^. We find that the amplitude of complex spike elicited calcium transients imaged from PC dendrites is indeed reduced during both NREM and REM sleep. This suggests that perhaps the number of spikelets riding on the climbing fiber induced complex spikes is reduced during sleep. There is only limited electrophysiological evidence for this in vivo. Mano (1970) reported that the number of spikelets in a complex spike waveform was reduced only during REM sleep with a similar distribution in the number of spikelets during wakefulness and slow wave sleep ^75^. Najafi and colleagues (2014) proposed two explanations for an increased amplitude of sensory evoked calcium transients ^71^. They suggest a small component of the increase could be attributed to a route other than the complex spike itself, most likely by way of parallel fiber to Purkinje cell inputs. However, for stronger sensory stimulation this contribution was deemed insufficient and that more likely an additional complex spike spikelet could contribute to the observed increase in calcium transient amplitude. Indeed, previous work supports a role for spikelets in driving additional calcium influx associated with complex spikes whose number was dependent on inferior olive oscillation phase ^82^. Both the phase and amplitude ^83^ of inferior olive subthreshold oscillations have been suggested to affect spikelet number. Given that we see a reduced calcium transient amplitude throughout sleep and not just during late NREM when cortical activity is likely to be partially gated ^84^, a reduced excitability of inferior olive neurons could be an additional contributing factor to a reduced number of spikelets and reduced calcium transients amplitudes across all sleep stages. Such reductions may be significant for memory consolidation during sleep since the induction of plasticity is affected by the number of spikelets^82^.

### Attenuation of the rate of complex spike co-activation during sleep

With our imaging approach it was possible to examine the level of co-activation of Purkinje cell dendrites within a cerebellar microzone. In our current analysis, we looked at the rate of occurrence of co-activations where at least 10% of Purkinje cell dendrites participated. Recent work suggests that the inferior olive in awake mice displays quasiperiodic rhythms in response to sensory input ^85^, which are relatively short lasting. Thus, if the level of cortical and midbrain input to the inferior olive is reduced during specific stages of sleep, one could also expect an impact on the level of synchronization of inferior olive oscillations and synchronous firing. We found that the co-activation rate was significantly greater during wakefulness even though synchronous firing still occurred during sleep. Synchronized firing in the inferior olive during sleep is likely to occur at specific time intervals during which cortical drive is strongly elevated such as at the cusp of synchronized cortical oscillations, or during eye movement or twitch related bursts of activity that occur during desynchronized or REM sleep ^86,87^

### Cerebellar sleep spindles gate cerebellar to cortical information transfer

During NREM sleep mice show cerebellar spindles and slow waves with directed coherence, providing evidence that during spindle frequencies information is transferred from the cerebellum to the cortex. This has previously been suggested to also occur in non-human primates ^88^. Whether cerebellar spindles play a role in gating out afferent sensory transmission to the neocortex is unknown ^26,54^. We found that twitch-like nuchal movements correlate with cerebellar spindles and muscle twitches triggered complex spike-evoked calcium transients both during wakefulness and sleep. In principle the olivary system is able to generate spindle frequency oscillations through a reverberating loop from the inferior olive, to the cortex, to the cerebellar nuclei and back to the inferior olive ^74,89^. However, we did not find reverberating complex spike activity with spindle frequencies during NREM sleep. Instead, our findings mirror recordings of the inferior olive in juvenile rats where activity increased sharply at the onset of nuchal activity during both wakefulness and sleep ^90^. The authors proposed that the twitches are accompanied by a corollary discharge that is relayed to the cerebellar cortex guiding predictions on the sensory consequences of movements ^91^. Previous work also found strong modulation of complex spike activity during REM sleep in juvenile rats ^92^. Twitching movements persist in the adult during sleep and thus may also contribute in adults to motor learning refinements ^93^. Studies in adult rats reported mostly ocular and masseter muscle rather than neck muscle activity during REM sleep ^94^. Implanting electrodes in the musculus rectus superior and musculus rectus lateralis could ease the association of REM state related movements with complex spike activity ^95^. One distinguishing feature in our recordings during REM sleep were two clearly discernible peaks in the triggered calcium response. Reverberation after a volley in the olivocerebellar loop could possibly contribute to such a secondary peak during REM sleep ^74,96^. The fact that throughout species, twitches correlate with spindles and cerebellar spindles with cortical spindles suggests a functional role for twitches and cerebellar spindles in cerebellar to cortical communication, and possibly plasticity as well as cerebellum-dependent memory ^27,56,66,69,93,97–100^. That twitches correlate with spindle activity may suggest a role for twitches in the process. To uncover how twitch reduction or an increase in twitches adjusts cerebellum-dependent motor memory processing is an important next step that needs to be solved before we can address the questions to what extent and how sleep may contribute to cerebellar processing.

## Acknowledgements

We are grateful to the following people for their contribution to the study: B. Winkelman, A. Court, L. Witter, and the whole CCC group for their contribution to useful discussions. This study was enabled by funding from the Netherlands Organization for Scientific Research (NWO-ALW 824.02.001; C.I.D.Z., NWO STEM - VBT 2021 19224 and NWO 863.14.005; C.B.C.), the Dutch Organization for Medical Sciences (ZonMW 91120067; C.I.D.Z.), Medical Neuro-Delta (MD 01092019-31082023; C.I.D.Z.), INTENSE LSH-NWO (TTW/00798883; C.I.D.Z.), ERC-adv (GA-294775 C.I.D.Z.) and ERC-POC (nrs. 737619 and 768914; C.I.D.Z.), the NIN Vriendenfonds for Albinism (C.I.D.Z.), the Dutch NWO Gravitation Program, and the Dutch Brain Interface Initiative (DBI2; C.I.D.Z.).

## Author contributions

C.B.C. and C.I.D.Z. conceived the project and designed the experiments. C.B.C, S.B. and P.M. analyzed the physiology data. C.B.C. and P.M. performed the awake extracellular recordings, C.B.C. performed the polysomnographic recordings, C.B.C and T.M.H performed the miniscope recordings, H.H.H and C.G performed surgeries for miniscope recordings, T.M.H analyzed the miniscope recordings, C.B.C. and S.L analyzed sleep recordings.

C.B.C., S.B., T.M.H., C.I.D.Z. wrote and edited the manuscript. C.B.C., C.I.D.Z. provided supervision.

C.I.D.Z. and C.B.C. acquired the funding.

## Methods

All protocols have been approved by the Animal Experiment committee of the Royal Netherlands Academy of Arts and Sciences (DEC-KNAW), and all procedures fulfil the European guidelines for the care and use of laboratory animals (Council Directive 86/6009/EEC). The mice had *ad libitum* access to food and water in their cages. They were housed in a stable at a temperature of 20±2°C, 60±20% humidity and with a timed 12-12 hours light-dark schedule. Experiments were performed during the light period, when mice were less active. Bedding material was provided for nest building.

### Surgical methods

#### Electrode surgery for polysomnographic recordings

C57BL/6 mice were used to obtain 4-24 hours polysomnographic recordings. All animals received Metacam (AUV 2 mg/kg) as analgesia at the beginning of the surgery. Eyes were protected and kept moist using eye drops drops (Duodrops, Ceva Santé Animale). Animals were anesthetized with isoflurane (4-5% induction, 0,8-1.5% maintenance; 0.2L/min O_2_ and 0.2 L/min air) and the head of the anesthetized mouse was placed in a stereotactic frame (David Kopf Instruments). During the whole procedure the body temperature of the mice was kept at 37°C using a feedback-controlled heating pad. The animal was shaved down to its shoulders, xylocaine was sprayed (10% Xylocaine, AstraZeneca) as a local analgesic, and the skin of the head was cut sagittally and then moved to the sides for exposure of the bone. The bone was etched with a phosphoric acid gel (37.5%, Kerr, Italy) and quickly washed with saline if needed. After drilling of holes for electrode implantation, primer (Optibond, Kerr) was applied on the bone and treated with UV light. Two holes were drilled (Foredom Drill K1070-2E, Blackstone Industries) for the bilateral implantation of M1 motor cortex EEGs (MC) (1.5 mm anterior and ± 2 mm lateral from *bregma*). We also drilled a whole and placed an electrode frontal as a reference, and on the parietal hemisphere very posterior. Additionally, two extra holes were drilled in order to reach the hippocampus (2 mm posterior and 1.5 mm lateral from *bregma*, and 0.2 mm deep from *duramatter*) and the pontine nuclei (3.8 mm posterior and 0.2 mm lateral to *bregma* and 5.8 mm deep from *duramatter*) for later LFP implantation; an EOG electrode was also implanted on their left eye muscle. Finally, an EMG was attached to the lateral neck muscle and another one was placed under the skin as a reference. Next, the three neck muscle layers were cut and moved to the sides in order to expose the occipital bone, without detaching the neck muscle EMG. A hole was drilled in the left hemisphere of the occipital bone to reach the cerebellum and an EEG implanted on it. EEGs, EMGs and EOGs were made from chloride silver wires, quadruple PTFE insulated for the EMGs, and soldered to IC connectors. All of them were bent before implantation to avoid the duramatter from punctuating. For a much easier implantation in deeper brain areas, the LFPs were made of insulated tungsten fibres, also soldered to IC connectors. Dental cement (Flowlime, Heraeus, Germany) was applied to the skull according to manufacturer’s specification (Kerr, Orange, California) to incorporate all the EEG, EMG, EOG and LFP electrodes as a solid block on top of the mouse’s skull. Post-operative analgesia was given subcutaneously with Meloxicam (2mg/kg Metacam, Boehringer Ingelheim Animal Health) and mice were placed under infrared light while recovering from anesthesia. Mice recovered for at least two days following surgery before subsequent experiments.

#### Craniotomy surgery for in-vivo electrophysiology

C57BL/6 mice were used for electrophysiological juxtacellular recordings of the cerebellum during physiological sleep. All animals received Metacam (AUV 2 mg/kg) as analgesia at the beginning of the surgery and were sprayed xylocaine (10% Xylocaine, AstraZeneca) in their neck muscles as a local analgesic. All animals followed almost an identical surgery procedure as previously explained for the electrode surgery. In N = 8 of these mice, no hippocampal or pontine LFPs nor the cerebellar EEG were implanted. For the remaining N = 9, a cerebellar EEG was implanted in the left hemisphere of the occipital bone; hippocampal and pontine LFPs were attached only in N = 5 out of these 9 mice. Additionally, a pedestal made of two soldered M1.4 nuts was attached to the head of all mice and cemented as a solid block to the electrodes with Primer (Optibond All-in-on, Kerr, Italy) and dental cement (Flowlime, Heraeus, Germany). This way the mouse’s head could be head-fixated with two screws during experiments and habituation. After the dissection of the three neck muscle layers and exposition of the occipital bone, a small window was also created over the skull. The location depended on the area of interest. Position of craniotomy for cerebellar nuclei was above Crus II to access the CN in a 43° angle. A thin layer of UV-curing primer (Optibond All-In-One, Kerr) was applied around the craniotomies and a recording bath was made using dental acrylic (Flowline and Charisma, Heraeus Kulzer). The recording bath was filled with 0.9% NaCl solution to keep the dura moist. Skin and muscles surrounding the bath were attached to the dental cement using tissue glue (Histoacryl, Aesculap). The dura in the craniotomy was carefully removed using a needle tip (30G x ½ inch, Microlance, BD). Directly after removing the dura, the recording bath was cleaned with 0.9% NaCl solution and filled with a low viscosity silicone elastomer sealant (Kwik-cast, World Precision Instruments) to protect the brain tissue from the air. Mice were given post-operative analgesia subcutaneously (Metacam 2 mg/kg) and were placed under an infrared light during recovery. Between recording sessions, the craniotomy was kept covered with silicone.

The orexin/EGFP mice followed a very similar surgery procedure to that of the C57BL/6 electrophysiology mice. All mentioned EEGs, EMGs and LFPs were implanted on them. Additionally, two extra holes were drilled over the lateral hypothalamic area (LHA, 1.7 mm posterior and 1 mm lateral from *bregma*) of these mice for future targeting of orexin neurons. Two plastic optic fibres (0.2 mm diameter, Thorlabs) were bilaterally implanted through these holes with a depth of 4 mm from *duramatter*, just 1 mm above from the LHA. The implants were fixed into place using dental cement (Super Bond, C&B).

#### Preparatory surgery in vivo calcium imaging

Mice were housed socially in groups of up to five mice prior to GRIN lens implantations, after which they were housed solitary. Around thirty minutes prior to surgeries mice received a first injection of analgesics. Directly prior to surgeries mice were anesthetized with 3% Isoflurane before being transferred to a stereotactic apparatus after which anesthesia was maintained at 1.1%-1.5% Isoflurane (flow rate: 0.3 ml/min O_2_). A small incision was made in the skin after hair removal and disinfection of the skin with iodine solution (5%) and alcohol (70%). Lidocaine (100 mg/ml, Astra Zeneca, UK) was then applied to the exposed skull and the periosteum removed. The center coordinates for GRIN lens placement were located and a small ink dot was placed at the correct location relative to bregma (cerebellar Simplex lobule, AP: −5.8 mm ML: 2.2 mm; lobule VI, AP: −7.4 mm, ML: 0.0 mm; cortex, AP: 1.4 mm, ML: 1.5 mm). Coordinates were scaled relative to the mean bregma-lambda distance (of 4.21 mm) as specified in Paxinos mouse brain atlas. Prior to drilling of the bone, mice received i.p. Injections of 15% D-Mannitol in saline (0.55 ml/25gr) to aid diffusion of virus particles after virus injection. A 2 mm circular craniotomy was then drilled centred around the marked location. In between drilling the skull was kept moist with sterile saline. The skull flap and dura were then removed and virus (Cerebellum: AAV1.CAG.FLEX.GCaMP6f/AAV1.CMV.PI.Cre.rBG mixed 1:1 and diluted in saline 1:3; Cortex: AAV1.Syn.GCaMP6f.WPRE.SV40 diluted in saline 1:3, UPenn Vector Core) was injected at four locations. At each location 25 nl of virus was injected once at 350, twice at 300 and once at 250 µm depth at a rate of 25 nl/min with a Nanoject II Auto-Nanoliter Injector (Drummond Scientific Company, USA). The craniotomy was covered with gelfoam (Pfizer, USA) soaked in sterile saline (0.9% NaCl, B. Braun Medical Inc, USA). The GRIN lens was lowered using a vacuum holder placed in the stereotactic apparatus until the lens surface touched the brain and then lowered an additional 50 µm. The edges of the craniotomy were sealed with Kwik-Sil (WPI, USA). Dental cement (Super-Bond C and B, Sun Medical, Japan) was then applied around the lens to secure it in place. Kwik-Cast (WPI, USA) was used to cover and protect the lens. At the end of the surgery animals received an s.c. injection of buprenorphine. For imaging of the cerebral and cerebellar cortex, a GRIN objective lens (1.8 mm diameter, 0.25 pitch, 64–519, Edmund Optics) was placed on the brain surface.After 2-3 weeks the scope holder was place

### Polysomnographic recordings

*In-vivo* baseline polysomnographic recordings were performed in a freely movement environment or home cage for 2-24 hours, min. 3 days after surgery. No habituation or handling was needed to perform these experiments. Recording pins to record sleep were attached under gas anesthesia of max. 5 minutes. The pins attached to their heads were connected through metal pins to an adapted MEA 60m Multichannel systems (adaptedMEA60, Multichannelsystems, Reutlingen, Germany), and the measured signal was digitalized with the MC_Rack utility.

### In-vivo (juxtacellular) electrophysiological recordings

#### Handling

Prior to surgery mice were handled 5-8 times in 2 weeks to reduce stress. After surgery the animals were handled and habituated to reduce stress prior to experiments.

#### Habituation

After recovery from surgery (normally 3 days) the mice were habituated to the electrophysiological setup. The mice were temporally anesthetized with isoflurane (1.5% in 0.5 l/min O2 and 0.2 l/min air) and placed in the setup by attaching the head with the pedestal in a stereotactic holder, restraining the body of the animal in a tube or circled plastic box. Some of their bedding was kept inside this circled box for the mice to feel more comfortable and increase the chances of sleeping. The habituation lasted for 20-30 minutes the first two days and then between 1-4 hours. The mice underwent at least three habituation sessions before recordings were made.

#### Recordings

During days of the recordings, the mice were placed in the electrophysiological setup after short isoflurane (3% in 0.5 l/min O2 and 0.2 l/min air, 30 min. post-anesthesia recovery time) anesthesia and EEG and EMG leads were connected to the IC connectors on the skull of the mouse. The EEG/EMG leads were connected to an amplifier (adaptedMEA60, Multichannelsystems, Reutlingen, Germany). The silicone was removed from the recording chamber and juxtacellular recordings were made using glass electrodes (Harvard Apparatus, Holliston, Massachusetts). The resistance of the electrodes varied between 6 and 13 MΩ. The glass electrodes contained internal solution consisting of: (in mM): 10 KOH, 3.48 MgCl2, 4 NaCl, 129 K-Gluconate, 10 hepes, 17.5 glucose, 4 Na2ATP, and 0.4 Na3GTP. The setup was connected to a Multiclamp 700B amplifier (Axon Instruments, Molecular Devices, Sunnyvale, California). The signal was digitized at 50 KHz with a Digidata 1440 (Axon Instruments, Molecular Devices, Sunnyvale, California, United States). Glass electrodes were advanced through the cerebellar cortex and Purkinje cells were identified by the presence of complex spikes and a pause in simple spiking after occurrence of a complex spike. All other neurons were identified by their firing patterns. First, spontaneous activity was recorded for each neuron for at least 60 seconds to check whether data quality was good. After the neuron was recorded for as long as possible. In the Orexin-ArchT mice, orexin neurons in the lateral hypothalamic area were inhibited by stimulating archaerhodopsin with light. These neurons were continuously illuminated for 60 seconds through optical fibers from a custom made LED light driver (590nm). Extracellular recordings and EEG/EMG recordings were continued during optical stimulation until several minutes after to assess the effect of the stimulation. After everything was stimulated, the neuron was recorded for as long as possible. For some neurons we also identified the recording location by adding Evans blue to the intracellular solution. We injected the die with pressure after a successful recording.

### Miniscope calcium imaging

#### Sleep deprivation device

A rotating drum (∅ 39 cm, height 37 cm), divided into 4 semicircular compartments by stationary central walls (Technicoplast, Echirolles, France) developed at the University of Grenoble and assembled, adapted and optimized at the NIN ^101^ was used for sleep deprivation prior to recordings to increase the likelihood for sleeping during a recording sessions. We added circular openings in the walls and placed half-circular openings at the boundary between wall and platform to allow for social interaction between mice, each placed in one of the compartments of the device. Drum, wall and lids consist of Plexiglas. The bottom consists of aluminum diamond plate to prevent mice from sliding. The bottom was covered with sawdust from the home cages. Water and food were provided. The devices are driven by a computer-controlled motor (MACDO-B1, JVL, Birkerød, Denmark), which runs bi-directionally. The motor is connected to the drum via a belt. During the sleep deprivation the device rotates 1 minute clock-wise, 1 minute counter-clockwise with a break of 10 seconds in between rotations at a speed of 2 rotations per minute.

Recordings from cerebellum and cerebral cortex were performed in the mouse home cage with bedding material, food and water *ad libitum*. Water was provided via a gel. The miniscopes were attached following brief anesthesia (3% isoflurane induction, 30 min. post-anesthesia recovery time) to allow mounting of one or two scopes and focal adjustment and sleep recording electrodes.

#### Data analysis sleep scoring

Sleep scoring was performed using the free access python3.6 package Wonambi 4.12 (GPLv3 license). Two independent raters scored the sleeping stages by eye, using 4 second epochs and differentiating between four sleeping states: Wake, NREM1/2, NREM3 and REM and a fifth “Undefined/Artefact” state for uncertainty. EEGs and LFPs were band-passed filtered between 0.5-30 Hz, but no filter was used for the EMGs or EOGs. All channels were referenced to an electrode that was placed on top of the brain in the frontal or parietal region or under the neck skin of the mouse (later shown as EMG reference). A 4 second epoch was rated as “Wake” whenever there was high activity in the EMG neck muscle channel. If such activity was shorter than one second, and surrounded by low EMG activity, this was rated as tiwtch or short awakening, and the scoring that followed was based on the other channels. “NREM1/2” was rated when K-complexes and/or spindles showed in any of the EEG motor cortex channels and the EMG activity was low, or whenever the EMG activity was low and the signal from the other channels didn’t fit with NREM3 or REM stages. The appearance of delta activity (1-5 Hz) in the motor cortices EEGs, along with low EMG activity, was rated as “NREM3”. Lastly, in order to score as “REM”, theta activity had to be observed in the hippocampus LFP, along with lower EMG activity than during the other sleep stages. Rapid eyemovements were also visible in the beginning of REM in the EOG channel but were not used as a requirement for REM scoring. The off-line analyses were performed using Clampfit (Axon Instruments, pCLAMP10.2) and MATLAB (Mathworks, Nantucket). LFP signals were down-sampled. Statistical analyses used are indicated in the text. Data normally show the mean ± SEM.

#### Data analysis calcium imaging

Timestamps for all scopes were logged to disk and in case of dual scope recordings, frames of the videos were aligned prior to motion correction using NoRMCorre ^73,102^ and signal extraction using CNMF-E. We used the following parameters in CNMF-E to segment of Purkinje cell dendrites and cerebral cortical neurons respectively (Purkinje cell dendrites: gSig = 7, gSiz = 40; cerebral cortical neurons: gSig = 7, gSiz = 10, for both merge_thr = [1e-1, 0.85, 0.5]). In the cerebellar stimulation experiments, neurons in cortex were classified as responders if the post-stimulus signal rose above the pre-stimulus mean+2σ. Onset times of these calcium transients were determined by fitting a sigmoid function to the transients and onset time was set to where the fitted function rose above mean+σ of the pre-stimulus baseline. For every cell, activity pre-stimulus and during stimulation was averaged per trial and statistically tested using paired t-tests. Fiji ^103^ was used for raw data inspection and to create videos. Analyses were performed in Matlab (Mathworks, Nantucket), Python 3.7 ^104^ and ^105^.

### Perfusion and histology

Mice were transcardially perfused with 4% PFA in 1x PBS (10 mM PO43−, 137 mM NaCl, 2.7 mM KCl). After fixation, brains were removed and kept overnight in 10% sucrose in PBS after which they were subsequently embedded in gelatin and left overnight in 30% sucrose in PBS. Brain sections (50 µm) were cut in a cryostat (LEICA CM3050 S), stained with DAPI and then mounted (Dako Fluorescence Mounting Medium, S3023). Confocal images were collected on a SP8 confocal microscope (Leica, Germany) and whole brain images were obtained by stitching multiple (1024 × 1024 pixel) acquisitions into a final image.

